# ZNK1 senses zinc and degrades zinc transporter mRNA in trypanosomes

**DOI:** 10.64898/2026.01.27.702079

**Authors:** Teresa Leão, Anna Trenaman, Michele Tinti, Gustavo Bravo Ruiz, Idálio J. Viegas, Luisa M. Figueiredo, Margarida Duarte, Ana Maria Tomás, David Horn

## Abstract

Like other cells, parasitic and other trypanosomatids sense and regulate Zn^2+^ transport, but the mechanisms involved remained unknown. Here we identify a trypanosome RNA-binding protein which specifically eliminates ZIP3 Zn^2+^-transporter mRNA in Zn^2+^-replete conditions. We first demonstrated that *Trypanosoma brucei ZIP3* mRNA abundance is subject to 3’-untranslated region (3’-UTR) and Zn^2+^-dependent negative control. A genome-wide RNA interference library screen, using a reporter associated with the *ZIP3* 3’-UTR, identified Tb927.11.9510 as a candidate Zn^2+^-sensor. We name this protein Zinc Nuclear Knuckles 1 (ZNK1) since it localises to the nucleus and contains several Zn^2+^-knuckle motifs. ZNK1 is conserved among trypanosomatids, and a PIN-domain suggests a ribonuclease-based mechanism. We validate ZNK1 as a *ZIP3* 3’-UTR dependent negative regulator and identify a GU-repeat motif in the *ZIP3* 3’-UTR that is predictive of negative control by ZNK1. We use Cas9-editing to knockout ZNK1, and RNA-seq to assess the consequences, revealing highly specific accumulation of *ZIP3* transcripts in *znk1*-null cells. We conclude that ZNK1 senses Zn^2+^-abundance and eliminates ZIP3 mRNA in a Zn^2+^-dependent manner. We suggest that trypanosomatid ZNK1 is an RNA-specific zinc finger nuclease that binds *ZIP3* 3’-UTRs and degrades *ZIP3* mRNA only when the tandem sensor modules are coordinated with Zn^2+^.

**Key points:**

- Trypanosome Zinc Nuclear Knuckles 1 (ZNK1) is a zinc-sensor that eliminates zinc transporter mRNA.
- ZNK1 negative control operates via the transporter mRNA 3’-untranslated region.
- The findings indicate that trypanosomatid ZNK1 is a conserved RNA-specific zinc finger nuclease.

## Introduction

Parasitic protozoa such as trypanosomatids face constant challenges in acquiring essential metabolites from their environments, as they navigate distinct host niches with varying nutrient profiles. *Trypanosoma brucei*, for example, is transmitted by tsetse flies and causes African trypanosomiasis in humans and livestock. As a result, these pathogens have evolved dedicated systems to sense fluctuations in nutrient and co-factor availability and to modulate acquisition and utilization. Because these parasites employ widespread polycistronic transcription, they depend almost exclusively on post-transcriptional control mechanisms, primarily mediated by RNA-binding proteins (RBPs), to reprogram gene expression and for environmental adaptation [1]. Understanding how parasites detect and respond to these nutritional cues is critical not only for uncovering basic aspects of their biology, but also for identifying new targets for therapeutic intervention.

Zinc (Zn^2+^) is an essential micronutrient required for the structural and catalytic functions of numerous proteins involved in a wide range of cellular processes. Yet, because excess intracellular Zn^2+^ is cytotoxic, primarily due to protein mismetallation [2–4], cells must tightly regulate acquisition, distribution, and storage. This is typically achieved through the coordinated action of membrane transporters (cellular and organellar importers and exporters), metal-binding proteins, and regulatory factors [5].

Despite the importance of zinc homeostasis for cell viability, the molecular mechanisms by which trypanosomatids sense and regulate zinc availability remain largely unexplored. In many systems, zinc is carried into cells by members of the ZRT/IRT-like protein (ZIP) family of transporters. These proteins translocate zinc and other metals from the extracellular environment or from intracellular compartments into the cytosol [6,7]. To date, only one ZIP has been characterised as a zinc transporter in trypanosomatids, the *Leishmania infantum* high-affinity zinc importer *Li*ZIP3 [8]. Genome annotations predict the presence of ZIP family members in *T. brucei* [9], including orthologues of *Li*ZIP3, but little is known about their regulation. Indeed, factors that mediate zinc-responsive gene expression have not been identified in any trypanosomatid.

Here we dissect how *T. brucei* coordinates the expression of zinc transporter genes in response to changes in zinc availability. We first characterise the regulation of ZIP3, a predicted ZIP-family zinc importer, and find that its mRNA is subject to post-transcriptional repression under zinc-replete conditions, by a mechanism involving the *ZIP3* 3’-UTR. We then devised a genome-wide RNA interference library screen [10] using a zinc-responsive reporter under the control of the *ZIP3* 3’-UTR. We identify multiple ZIP-family members, including ZIP3, as effectors of zinc transport and a putative regulatory RNA-binding protein, which we named Zinc Nuclear Knuckles 1 (ZNK1); the protein is nuclear-localised and contains several Zn^2+^-knuckle motifs and a putative ribonuclease domain. We identify a GU-repeat motif in the *ZIP3* 3’-UTR that is predictive of negative control by ZNK1 and show that ZNK1 is indeed a zinc sensor and negative regulator of *ZIP3* expression.

Specifically, CRISPR-Cas9-mediated knockout of ZNK1 led to specific accumulation of *ZIP3* transcripts in zinc-replete conditions, suggesting that ZNK1 mediates zinc-dependent mRNA degradation. Together, our results reveal a novel post-transcriptional regulatory mechanism by which trypanosomes maintain zinc homeostasis. Thus, ZNK1 is a zinc-sensor and highly selective and specific effector of zinc-transporter mRNA turnover.

## Results

### *T. brucei ZIP3* mRNA is subject to zinc- and 3’-UTR-dependent turnover

The *Leishmania infantum* zinc importer *Li*ZIP3 is negatively regulated at the mRNA level in response to zinc [8] and we reasoned that the *T. brucei* orthologue would be similarly regulated. The ZIP3 orthologue in *T. brucei* is encoded by a syntenic tandem cluster on chromosome 11 (Tb927.11.9000 – Tb927.11.9030), which is preceded by a diverged paralogue *ZIP3’*, Tb927.11.8990. The *T. brucei ZIP3* genes encode proteins with ≈89 to 98% similarity and share similar 3’-UTRs, while the *ZIP3’* gene has a distinct 3’-UTR. Notably, the *ZIP3* 3’-UTR registered negative regulatory function in a recent massive parallel reporter assay [11].

Our first step was to determine whether *ZIP3* or *ZIP3’* expression responded to zinc or iron, two of the most common metals transported by ZIP family members [12–14]. Bloodstream form (BSF) *T. brucei* were grown for 1-3 h in zinc-sufficient (ctrl), zinc/iron-surplus (Zn/Fe) or zinc/iron-limited medium generated by treatment with the chelators TPEN (for zinc) and DFO (for iron), and expression of *ZIP3* and *ZIP3’* mRNA was then assessed using RT-qPCR. *ZIP3* displayed robust and significant induction in zinc-limited medium, with transcript levels increased >4-fold relative to growth in control or zinc-surplus medium (Fig. 1A). This response was zinc-specific, since only addition of zinc, but not iron, abrogated TPEN-induced upregulation (TPEN+Zn), while supplementation with iron had no significant effect (Fig. 1A). In similar assays, *ZIP3’* showed a significant, but relatively moderate response in TPEN+Fe (Fig. 1B).

**Figure 1.**
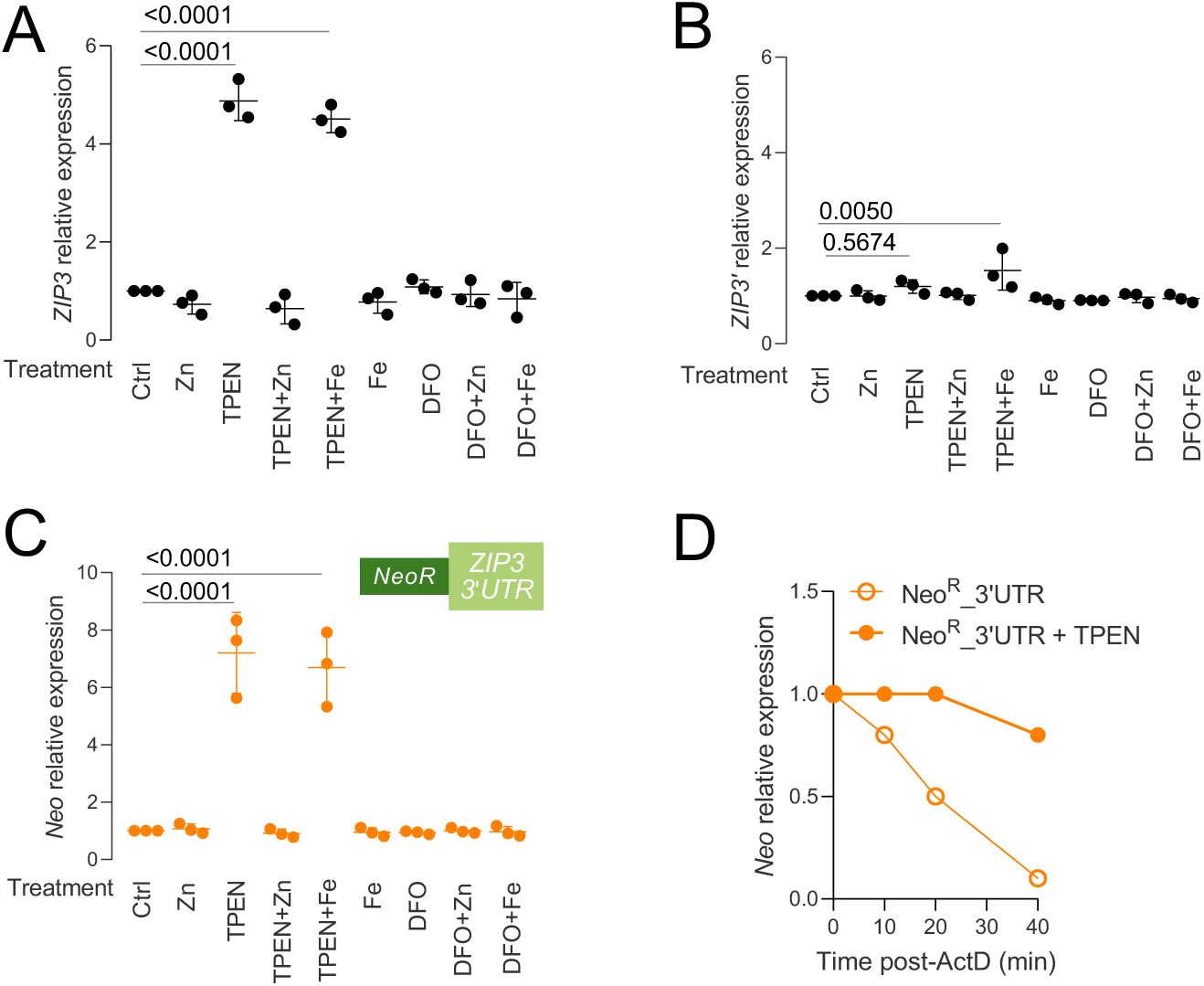
*T. brucei ZIP3* mRNA is subject to zinc- and 3’-UTR dependent turnover. (**A-B**) RT-qPCR assays were used to assess the abundance of *ZIP3* (**A**) or *ZIP3’* (**B**) transcripts in *T. brucei* grown in zinc-sufficient (ctrl), zinc/iron-surplus (Zn/Fe, 100 µM, 1 h) or zinc/iron-limited medium; TPEN (for zinc, 10 µM, 2 h) and DFO (for iron, 10 µM, 2 h). Data represents fold change in expression relative to the “ctrl”, normalised to β-tubulin transcript abundance. *p*-values were derived using one-way ANOVA tests with Sidak multiple comparisons. Mean ± SD from three independent experiments are shown. (**C**) RT-qPCR assays as in A-B but to assess the abundance of the Neo^R^ reporter transcript under the control of the *ZIP3* 3*’*-UTR. (**D**) RT-qPCR assays were used to assess the Neo^R^ *ZIP3* 3*’*-UTR reporter transcript after inhibition of transcription with actinomycin D (ActD, 5 µg/mL), in control or TPEN-treated cells.

In trypanosomatids, elements regulating gene expression are often embedded within the 3’-UTR [11]. To determine whether this is the case for ZIP3, we generated a reporter strain expressing a G418 resistance gene (Neo^R^) under the control of the *ZIP3* 3’-UTR. To determine whether the 3’-UTR impacted Neo^R^ expression, we subjected the reporter strain to the same metal supplementation/limitation conditions detailed above. RT-qPCR analysis to assess Neo^R^ expression revealed a response to zinc that mirrored the *ZIP3* response, whereby Neo^R^ mRNA expression was significantly induced by TPEN, with this effect only reversed by the addition of zinc (Fig. 1C). Sensing through the ZIP3 3’-UTR thus enables *T. brucei* parasites to adapt to zinc availability.

To determine whether the *ZIP3* 3’-UTR drives mRNA turnover in the presence of zinc, we used actinomycin to inactivate transcription and again assessed reporter mRNA abundance in zinc replete or depleted medium. We observed that under zinc-limiting conditions (TPEN), the transcript was relatively stable, whereas in zinc-sufficient conditions the transcript decayed with a half-life of approx. 20 min (Fig. 1D). These results indicated that the *ZIP3* transcript was indeed turned over in the presence of zinc.

### A genome-wide screen for loss of *ZIP3* 3’-UTR-based negative control

We next sought to identify the regulatory factor or factors responsible for *ZIP3* mRNA turnover. We demonstrated above that a *Neo^R^*transcript under the control of the *ZIP3* 3’-UTR is unstable in zinc-replete medium and is stabilised when zinc is limiting. We, therefore, designed a genome-wide RNA interference (RNAi) target sequencing (RIT-seq) screen for knockdowns that increase Neo^R^ *ZIP3* 3’-UTR reporter expression in zinc-replete medium. Although genome-wide RIT-seq screens have been used widely for unbiased forward genetics and gene discovery, few screens have been used to identify factors operating via 3’-UTRs [15,16].

To establish the RNAi library, we first transformed 2T1-T7 cells [17] with the zinc-sensitive Neo^R^ *ZIP3* 3’-UTR reporter construct; these cells also express the tetracycline repressor and T7 RNA polymerase which allow for inducible expression of the RNAi library [10] (Fig. 2A). We then determined the effective concentration of G418 to inhibit the growth of these cells by 50 % (EC_50_) to be 4 µM (Supplementary Fig. 1), which reflects Neo^R^ expression in these cells. The Neo^R^ *ZIP3* 3’-UTR reporter strain was then used to generate the RNAi library [10].

**Figure 2.**
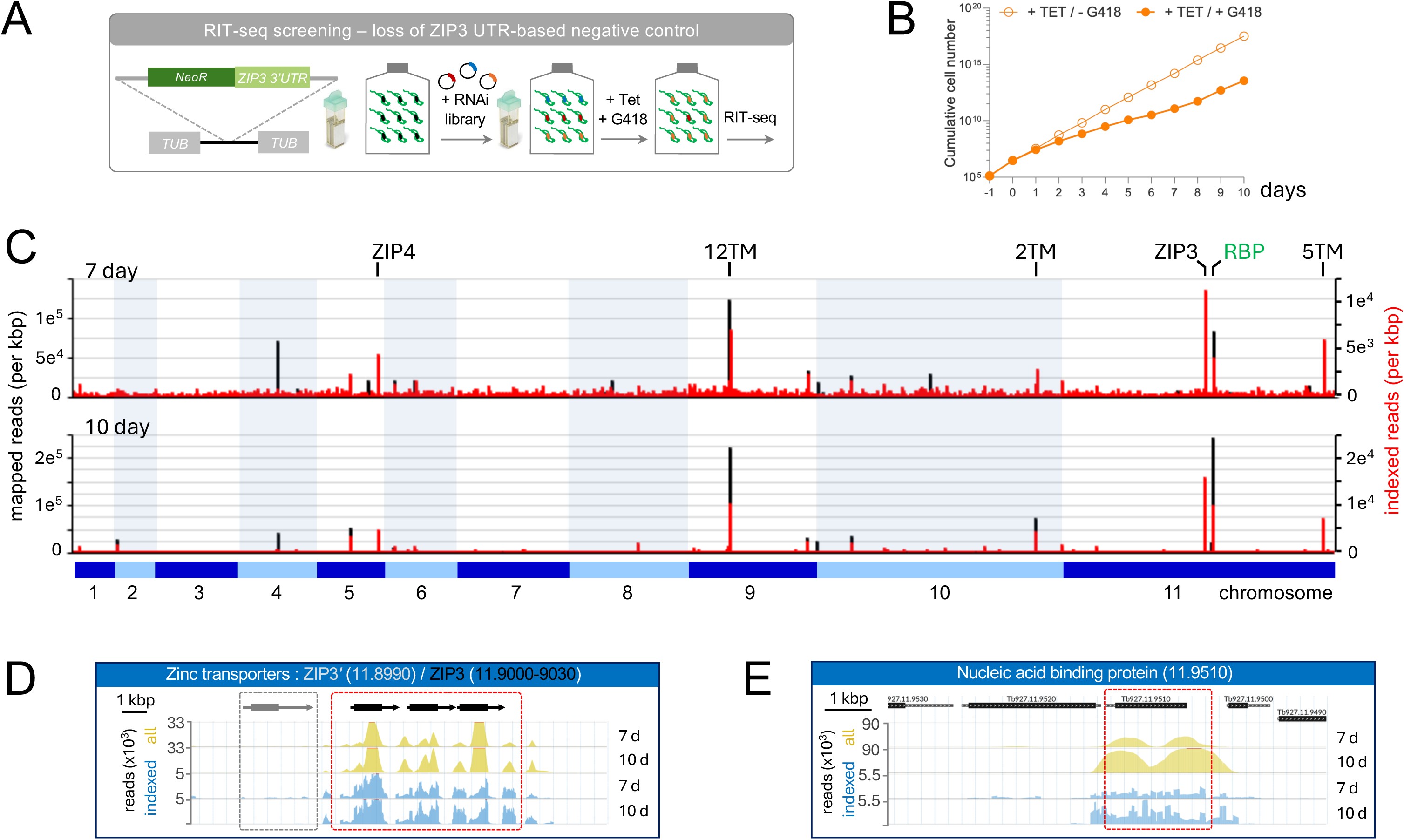
A genome-wide screen for loss of *ZIP3* 3*’*-UTR-based negative control. (**A**) Schematic representation of the genome-scale RNAi (RIT-seq) screen. *T. brucei* expressing the zinc-responsive Neo^R^ *ZIP3* 3*’*-UTR reporter were used to assemble an RNAi library. RNAi knockdown was induced with tetracycline (Tet, 1 µg/mL), G418 selection for loss of *ZIP3* 3*’*-UTR-based negative control was applied 24 h later, and genomic DNA was extracted for RIT-seq profiling. (**B**) Cell density was monitored during RNAi library selection with G418. Two parallel libraries were induced with tetracycline, one was selected with G418 and the other served as the control. DNA was collected from the G418-selected library after 7 and 10 days. (**C**) Genome-scale RIT-seq profiles show enriched RIT-seq fragments. Data for both day 7 (D7) and 10 (D10) are shown. Black peaks represent total mapped reads per kilobase, whereas red peaks represent reads that incorporate an index sequence from the RNAi plasmid library. (**D**) RIT-seq profiles show enriched RIT-seq fragments at the *ZIP3/3’* locus (*ZIP3*, red box; *ZIP3’*, grey box). Total reads (yellow) and indexed reads (blue) are shown for both D7 and D10. (**E**) RIT-seq profiles as in D but for the Tb927.11.9510 locus.

To run the screen, the RNAi library was induced with tetracycline and grown in the presence of G418 at 8 µM (2 × EC_50_). Cell density was monitored for ten days under these conditions and compared to a parallel control culture grown without G418 selection (Fig. 2B). G418 selection impaired library growth as expected, and we harvested cells after seven and ten days when G418 had reduced cell number >1300-fold and >8800-fold relative to the control culture (Fig. 2B). Genomic DNA was extracted from the harvested cells, and DNA libraries were generated by PCR amplification of the RNAi target fragments. The amplicons were deep sequenced, and reads were mapped to the *T. brucei* reference genome. The genome-scale maps reveal substantial enrichment of several RIT-seq fragments, validated by the presence of flanking index sequences present in the plasmid library (Fig. 2C, Supplementary Data 1). Notably, those enriched RIT-seq fragments were very similar after either seven or ten days of library selection, but with fewer background reads after ten days; we considered those top six hits, on chromosomes 5 and 9-11 that registered >3000 reads incorporating the index sequence from the RNAi plasmid library in both datasets (red bars in Fig. 2C).

Five of these hits identified genes with between two and twelve predicted transmembrane domains, including *ZIP3* (Fig. 2D, red box). ZIP3 knockdown likely depletes intracellular pools of zinc, essentially mimicking TPEN-treatment (see Fig. 1C) and this probable indirect activation of the *ZIP3* 3’-UTR reporter by ZIP3 knockdown provided excellent validation for our screen. The same rationale applied to Tb927.5.4140, which is also annotated as a ZIP-family member (Supplementary Fig. 2A), and now referred to as ZIP4; unlike *ZIP3*, *ZIP4* expression was not sensitive to zinc availability (Supplementary Fig. 2B). Notably, *ZIP3’* did not register as a hit in the screen (Fig. 2D, grey box), suggesting a distinct role for this paralogue. The remaining three transmembrane proteins included a Major Facilitator Superfamily (MFS) transporter (Tb927.9.6360-80) previously linked to aminoglycoside-resistance [18], suggesting a role in G418-resistance rather than zinc transport. Accordingly, we suspect that the remaining two predicted transmembrane proteins play roles in either zinc or G418 transport.

We were particularly interested in identifying the regulatory factor or factors responsible for *ZIP3* mRNA turnover, and the final hit in the RIT-seq screen (Tb927.11.9510) was annotated as a putative nucleic acid binding protein. This gene is on the same chromosome as the *ZIP3* array but these loci are in separate polycistrons and are separated by 116 kbp. Tb927.11.9510 presented an excellent candidate *ZIP3* regulator (Fig. 2E).

### Tb927.11.9510 / ZNK1 is a conserved probable zinc sensor

*In silico* analysis of the protein encoded by the *Tb927.11.9510* gene revealed seven zinc knuckle motifs (CxxCxxGHxxxxC) and one canonical zinc finger motif (CCHC), followed by a PIN (PilT N-Terminus) potential ribonuclease domain (Fig. 3A), features entirely consistent with a zinc-sensing RNA-binding protein that can degrade *ZIP3* mRNA. Indeed, ZNK1 is highly conserved among the parasitic trypanosomatids, *T. brucei*, *T. cruzi* and *L. infantum* (Fig. 3A), consistent with a conserved role in zinc-sensing and *ZIP3* mRNA turnover. Zinc knuckles and zinc fingers are small cysteine/histidine-rich modules that coordinate Zn^2+^ ions, and are typical of metal-dependent RNA-binding proteins (RBP); they link metal binding to conformational changes that stabilise the protein and/or facilitate RNA interaction [19,20]. These motifs are also conserved in number and spacing (Fig. 3A) and consensus sequence (Fig. 3B) in the ZNK1 orthologues. The PIN domain likely adopts an RNase-like fold, with conserved acidic residues coordinating a divalent metal ion essential for nucleic acid cleavage [21]. The protein encoded by *Tb927.11.9510* has been localised to the nucleus in insect-stage *T. brucei* [22] (and see below). We therefore named Tb927.11.9510 Zinc Nuclear Knuckles 1 or ZNK1.

**Figure 3.**
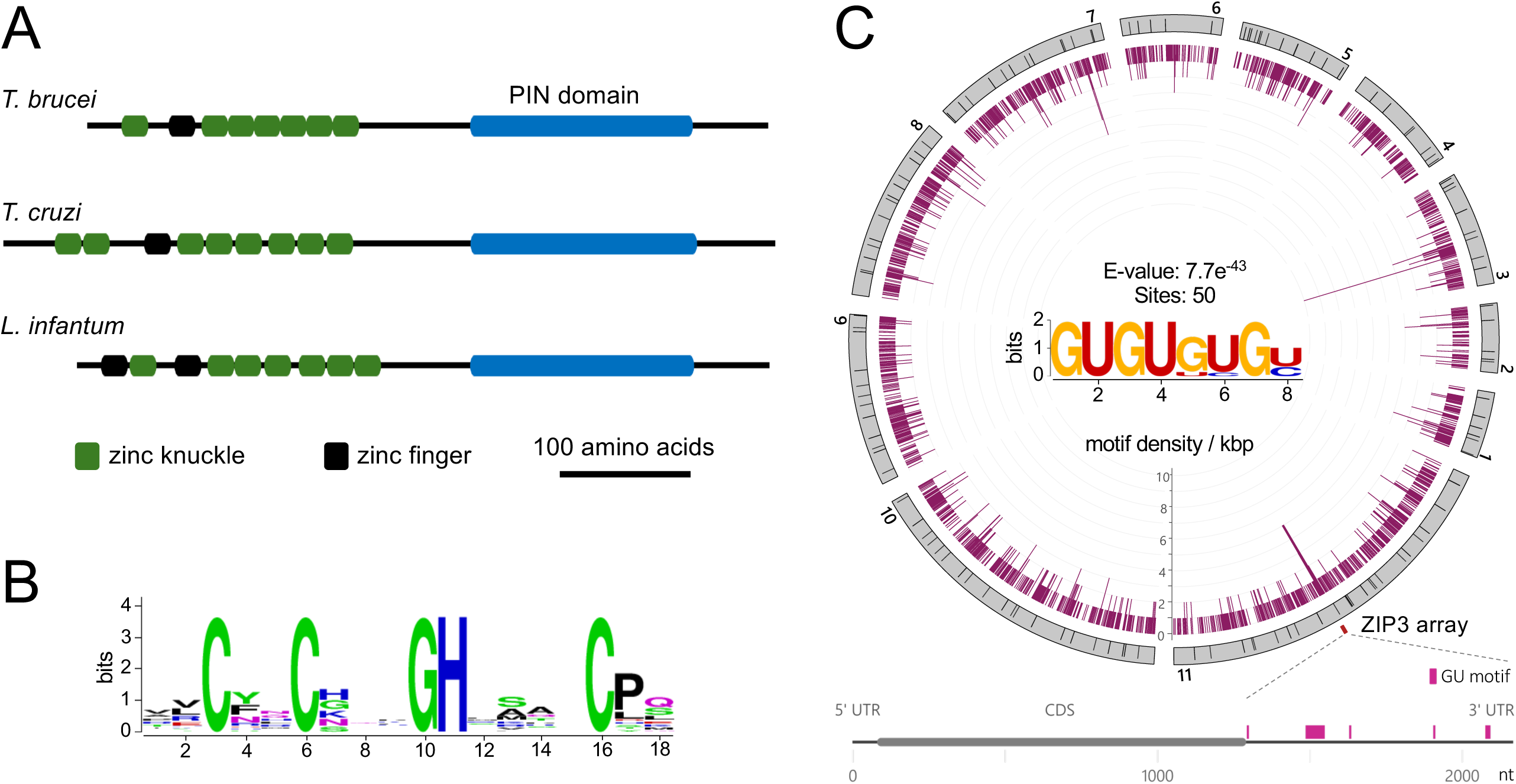
Tb927.11.9510 / ZNK1 is a conserved probable zinc sensor. (**A**) Schematic representation of the protein encoded by Tb927.11.9510, and the orthologues in *T. cruzi* (TcCLB.511589.110) and *Leishmania infantum* (LINF_360040200). Zinc knuckle motifs (CxxCxxxGHxxxxC, green), zinc finger motifs (CCHC, black) and the PIN putative nuclease domain (blue) are shown in each protein. (**B**) The sequence logo shows conserved residues in zinc knuckle motifs (n = 22) from the three proteins shown in A. (**C**) Motif enrichment analysis revealed a GU-repeat motif in *ZIP3* 3’-UTRs from multiple trypanosomatids. Density of this motif across the ‘427’ genome assembly is shown on the circular karyotype plot (inner circle). The outer circle shows chromosomes and promoters (black bars) for polycistronic transcription units. The lower panel shows localization of GU-repeat motifs in the *ZIP3* (Tb427_110096600) mRNA; the motif is only found in the 888-b 3’-UTR.

The conserved arrangement of zinc knuckle motifs in ZNK1 suggested a mechanism involving recruitment to *ZIP3* mRNA via binding to a repeated *cis*-acting element. To identify candidate *cis*-acting sequences, we searched for motifs that were over-represented in *ZIP3* 3’-UTRs from multiple trypanosomatids. This analysis revealed a GU-repeat motif at fifty sites in only eleven sequences (Fig. 3C, E value 7.7^−43^). Although there are many copies of this 8-b GU-repeat motif in the *T. brucei* genome, a genome-wide search revealed the highest concentration of this motif at the *ZIP3* locus (Fig. 3C), present at five locations within each 3’-UTR (Fig. 3C, lower panel); the latest annotation indicates thirteen tandem arrayed *ZIP3* genes (Tb427_110096600 – 110097800, https://tritrypdb.org). Such a repeated *cis*-acting sequence could act cooperatively to recruit a multi-zinc finger protein such as ZNK1.

### ZNK1 operates via the *ZIP3* 3’-UTR

To characterise ZNK1 in more detail, we first analysed ZNK1 transcript abundance in various conditions of zinc availability. Figure 4A shows that the *ZNK1* transcript is not substantially regulated by zinc concentration in the culture media. We next assembled a bloodstream-form *T. brucei* strain expressing N-terminally c-myc tagged ZNK1 (^myc^ZNK1) from its native locus. Immunofluorescence microscopy showed that ^myc^ZNK1 was strongly enriched in the bloodstream-form *T. brucei* nucleus (Fig. 4B). Neither ZNK1 subcellular localization (Fig. 4B) nor abundance (Fig. 4C) were substantially changed following zinc chelation with TPEN.

**Figure 4.**
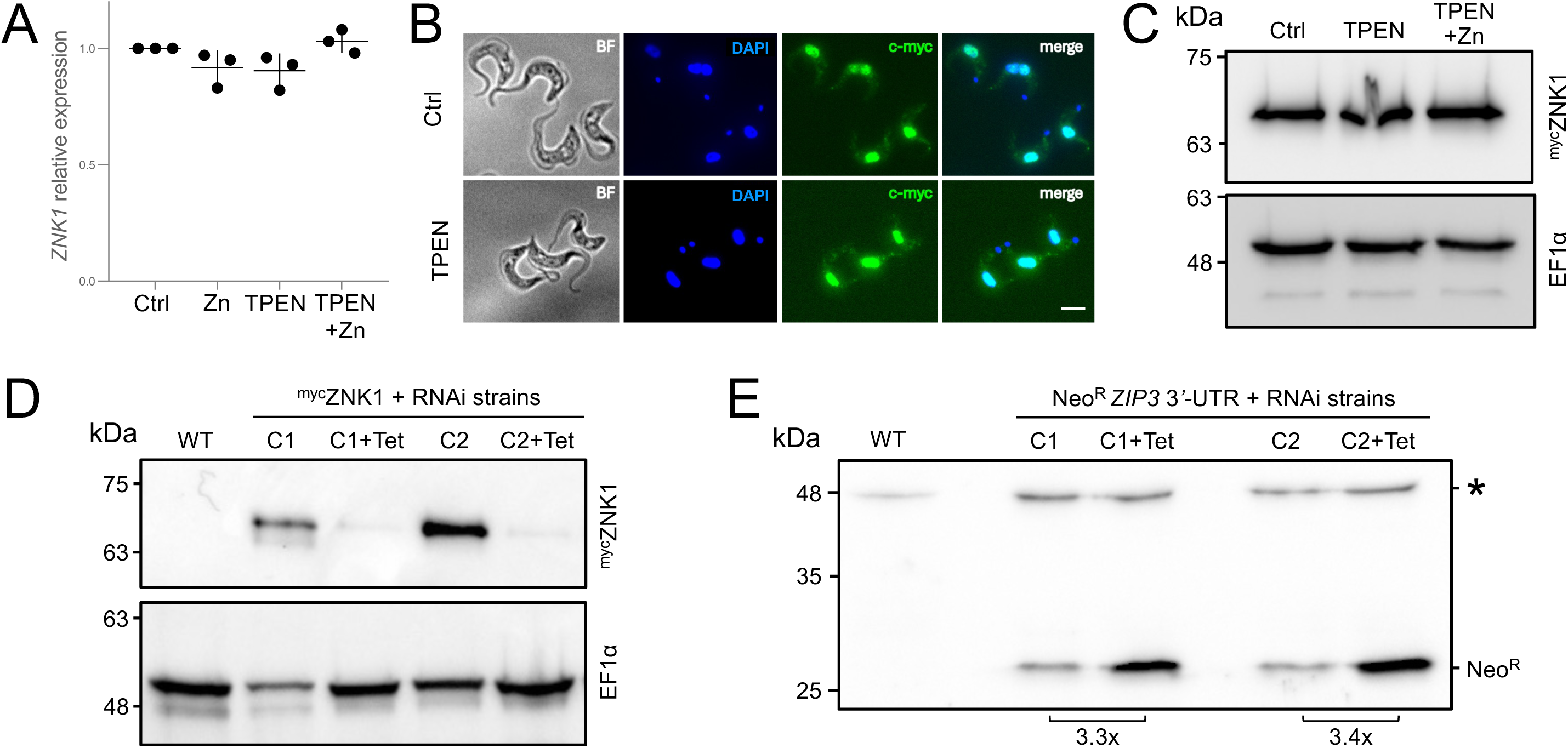
ZNK1 operates via the *ZIP3* 3*’*-UTR. (**A**) RT-qPCR assays were used to assess the abundance of *ZNK1* transcripts. Other details as in Fig. 1B. (**B**) Immunofluorescence microscopy showing the localization of ^myc^ZNK1 in control (ctrl) or zinc depleted (TPEN, 2 h) medium. Scale bar = 5 µm. (**C**) The protein blot shows the expression of ^myc^ZNK1 in control (ctrl), zinc depleted (TPEN, 10 µM, 2h) or zinc supplemented (TPEN+Zn, 10 µM + 100 µM, 2 h + 1 h) medium. The EF1α panel serves as a loading control. (**D**) The protein blot shows efficient ^myc^ZNK1 knockdown following induction of *ZNK1* RNAi using tetracycline (Tet) for 24 h. The EF1α panel serves as a loading control. WT, wild-type, C1-2, clones 1 and 2. (**E**) The protein blot shows expression of the Neo^R^ *ZIP3* 3*’*-UTR reporter before and after knockdown of *ZNK1* for 24 h. A non-specific band (*) serves as a loading control. Relative increase in the Neo^R^ signal following ZNK1 knockdown is indicated below the blot.

We next used ZNK1 knockdown to determine whether ZNK1 does indeed operate via the *ZIP3* 3’-UTR. A bespoke ZNK1 RNAi construct was used for robust knockdown of ^myc^ZNK1 (Fig. 4D) while ZNK1 knockdown using this same construct led to increased expression of the Neo^R^ *ZIP3* 3’-UTR reporter (Fig. 4E). This result phenocopied zinc limitation and was entirely consistent with identification of ZNK1 in the RIT-seq screen. Thus, ZNK1 mediates negative regulation via the *ZIP3* mRNA 3’-UTR when zinc availability is sufficient.

### Specific accumulation of *ZIP3* transcripts in *znk1*-null cells

To determine whether ZNK1 controls the abundance of *ZIP3* and/or other mRNAs, we performed CRISPR-Cas9-mediated knockout of the *ZNK1* gene in *T. brucei* bloodstream-form cells. From two independent transfections, we selected two *znk1* null clones where both *ZNK1* alleles were replaced with a Neo^R^ cassette, as determined by PCR (Supplementary Fig. 3), and genome sequencing was used to validate specific deletion of the *ZNK1* gene in both of these strains (Fig. 5A).

**Figure 5.**
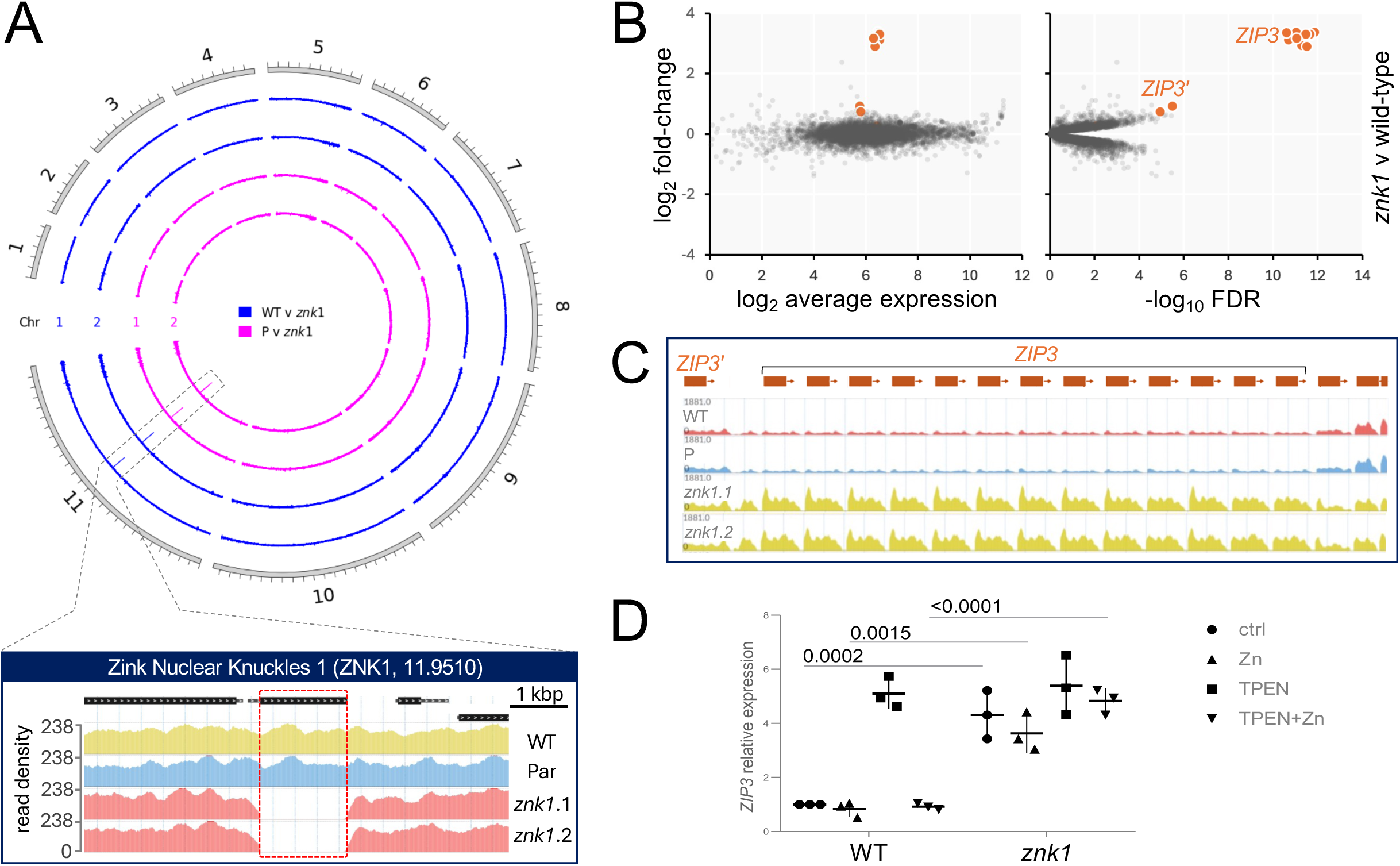
Specific accumulation of *ZIP3* transcripts in *znk1*-null cells. (**A**) The circular plot shows whole genome sequencing data for two independent *znk1* null strains compared to wild-type (WT) or parental (P) control strains, revealing specific knockout of the ZNK1 gene on chromosome 11. The lower, zoomed panel shows precise ZNK1 deletion. (**B**) RNA-seq analysis comparing *znk1* null and wild-type cells. *ZIP3* and *ZIP3’* transcripts are highlighted. The data represent averages from three independent technical replicates in each case. n = 8013. (**C**) RNA-seq analysis showing the *ZIP3*/*3’* locus (37 kbp) in the wild-type (WT) and parental (P) control strains and in the *znk1* null strains. (**D**) RT-qPCR assays were used to assess the abundance of *ZIP3* transcripts in wild-type (WT) and *znk1* null cells. Other details as in Fig. 1B.

To assess the impact of ZNK1 on mRNA abundance, we performed RNA sequencing (RNA-seq) under standard (zinc-sufficient) conditions with parental cells and both *znk1* null strains. RNA-seq confirmed *ZNK1* knockout with >4000 reads mapped to this gene in the parental strain and zero reads following knockout (Supplementary Data 1). RNA-seq also revealed a highly significant increase in *ZIP3* transcript abundance (log2 fold-change >3, False Discovery Rate [FDR] <1e^-11^) in the *znk1* null strains (Fig. 5B-C, Supplementary Data 1); *ZIP3’* transcript abundance was also significantly, although less substantially, increased (log2 fold-change 0.9, FDR <1e^-5^). We further validated increased *ZIP3* mRNA abundance in *znk1* null cells using RT-qPCR, showing that *ZIP3* transcripts were elevated in the absence of ZNK1 independent of zinc availability in the culture media (Fig. 5D). Using the same approach, we observed a moderate, although not significant, increase in the abundance of *ZIP3’* transcripts and no increase in the abundance of *ZIP4* transcripts (Supplementary Fig. 4).

Together, our results show that *ZIP3* transcripts are turned over in a 3’-UTR dependent and ZNK1-dependent manner when zinc supply is sufficient. Thus, we establish ZNK1 as a zinc sensor and post-transcriptional repressor of *ZIP3* transporter expression.

## Discussion

Regulation of zinc homeostasis is essential for cellular function and relies on the coordinated activity of zinc transporters and zinc storage [23]. However, the mechanisms by which zinc availability is sensed and translated into changes in transporter expression have remained incompletely characterised, in trypanosomatids and in many other cell types. Using 3′-UTR reporter assays, genome-wide RNA interference (RIT-Seq) screening [15,24], and transcriptomics, we identified and characterised *T. brucei* Zinc Nuclear Knuckles 1 (ZNK1), a zinc sensory RNA-binding protein required for zinc-dependent control of the *T. brucei* zinc transporter ZIP3. ZNK1 is highly conserved among the trypanosomatids and is therefore likely to be the key post-transcriptional zinc homeostasis regulator in these important parasites.

We first found that *T. brucei ZIP3* mRNA turnover is increased in the presence of zinc, and by a mechanism involving the 3′-UTR. Exploiting this information, we used a drug-resistance marker under the control of the *ZIP3* 3’-UTR as a reporter in a genome-wide RIT-seq screen. The screen identified zinc transporter knockdowns that reduced zinc availability, leading to activation of the reporter, and also identified the putative regulator, ZNK1. We validated ZNK1 in bespoke RNAi strains and also used CRISPR-Cas9 to knockout ZNK1; these cells were defective for zinc-dependent negative control of both *ZIP3* mRNA and the *ZIP3* 3’-UTR reporter.

Trypanosomatid ZNK1 incorporates seven or eight zinc knuckle motifs and one to two CCHC-type zinc finger motifs in the N-terminal region. These zinc-binding motifs are present in many eukaryotic RNA-binding proteins, and are suggestive of a role in zinc-sensing [25]. ]. Zinc knuckles are compact Cys/His-rich modules that typically engage single-stranded RNA, often acting as RNA “chaperones” that stabilise transient structures. When present in tandem, they can create an extended, modular binding surface that can recognise both sequence features and local RNA architecture along a 3′-UTR [26]. In trypanosomatid ZNK1, the presence of several knuckle motifs is therefore consistent with high-affinity and selective recognition of *ZIP3* 3′-UTRs. Because coordination of Zn^2+^ is integral to knuckle folding, changes in metal occupancy are expected to modulate the stability and orientation of the knuckles and, thus, RNA binding.

The C-terminal portion of ZNK1 encodes an NYN/PIN family, PIN superfamily, putative nuclease domain with conserved acidic residues typical of a metal-dependent catalytic center [21]. Other PIN-domain proteins in trypanosomatids include RRP44/Dis3 [27,28], protein*-*only RNase P enzymes [29], and ESB2 [30], which participate in rRNA/tRNA processing or subtelomeric expression-site-associated-gene control, respectively. ZNK1 is a unique member of the NYN/PIN family with an architecture suggestive of endoribonuclease activity coupled to a micronutrient-sensor domain. Thus, ZNK1 architecture is entirely consistent with our demonstration of zinc-dependent turnover of *ZIP3* mRNA. An Alpha Fold model [31] shows how ZNK1 may interact with ZIP3 mRNA (Fig. 6A).

**Figure 6.**
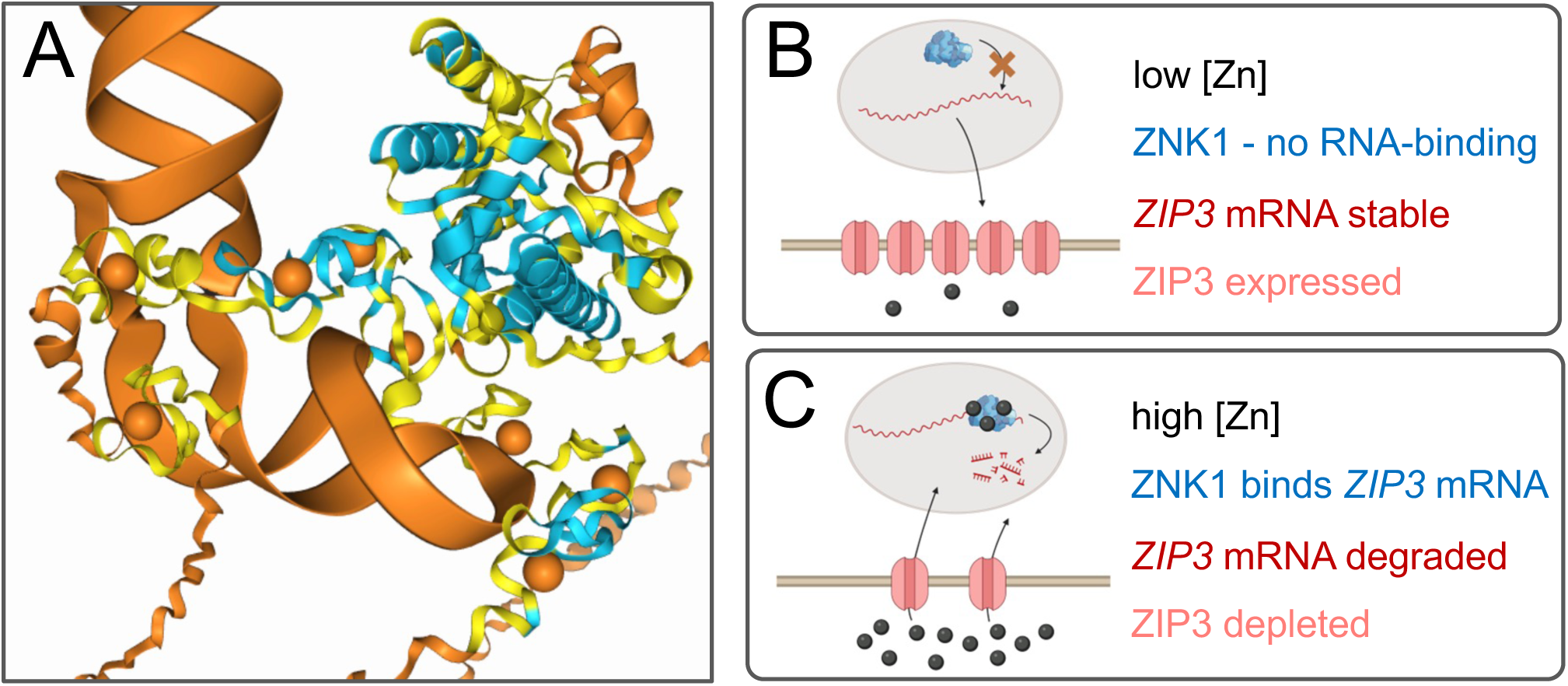
A model for zinc sensing and *ZIP3* mRNA turnover by ZNK1. (**A**) The AlphaFold model (https://alphafoldserver.com/) shows how ZNK1 may bind *ZIP3* mRNA, thereby recruiting the nuclease domain to its substrate. Eight Zn^2+^ ions (orange spheres) are included. Confidence: blue, confident; yellow, low; orange, very low. (**B**) The schematic model indicates how ZNK1 remains inactive in zinc-limiting environments, facilitating zinc uptake by ZIP3. (**C**) The schematic model indicates how ZNK1 binds and degrades *ZIP3* mRNA in zinc-sufficient environments.

*ZIP3* mRNA regulation by ZNK1 is remarkably specific. Nuclear localization of ZNK1 also suggests that *ZIP3* mRNA is turned over in the nucleus, possibly immediately following transcription. We suggest a model whereby under limiting zinc conditions, the zinc-knuckles in ZNK1 fail to fold, and fail to bind *ZIP3* mRNA which can then be effectively exported from the nucleus and translated, facilitating zinc uptake (Fig. 6B). When ZIP3 transporter activity provides a sufficient supply of zinc, the knuckles efficiently fold such that ZNK1 binds and turns over *ZIP3* mRNA (Fig. 6C). Indeed, we identify a strong candidate GT-repeat motif and *cis*-acting sequence likely required for productive ZNK1-binding. Thus, zinc-dependent changes in the RNA-binding module facilitate highly specific recruitment of this nuclease to target substrates. This simple sensing and feedback model likely explains zinc homeostasis control by the conserved ZNK1 RBP in the trypanosomatids.

The specificity of ZNK1 for ZIP3 contrasts with systems in which a single factor coordinates a broad zinc response, such as Zap1 in budding yeast, MTF-1 in mammals or the uptake regulator Zur in bacteria, which control zinc uptake and storage regulons as well as the expression of additional adaptive proteins at the transcriptional level [32–34]. In those cases, zinc availability is sensed by DNA-bound transcription factors. Thus, post-transcriptional control in *T. brucei* is distinct and facilitates nutrient-responsive control in the context of widespread polycistronic transcription. Similarly, the *T. brucei* NT8 nucleobase transporter mRNA is repressed by the PuREBP1/2 complex in a 3′-UTR PuRE stem–loop dependent manner when purines are abundant [33,35,36]. Similar principles have also been described for glucose- and amino acid–responsive post-transcriptional controls in *T. brucei* [37–39].

ZNK1 therefore extends an emerging paradigm in which trypanosomes implement nutrient sensing through RNA-centric mechanisms, whereby RBPs selectively bind regulatory 3′-UTRs, rather than employing classical transcription factor–based feedback loops.

In conclusion, our study reveals a zinc-dependent post-transcriptional mechanism that modulates zinc transporter expression in *T. brucei*. ZNK1 emerges as a key component of a zinc-responsive circuit linking metal availability to mRNA stability. This work uncovers a new dimension of parasite metal homeostasis and illustrates how early-branching eukaryotes can repurpose RNA decay machinery to achieve precise adaptation to fluctuating micronutrient availability. ZNK1 conservation suggests that this zinc sensor negatively controls zinc uptake to maintain appropriate zinc supply in the varying host environments encountered by multiple important parasitic trypanosomatids.

## Methods

### *T. brucei* growth and manipulation

Bloodstream form Lister 427 *T. brucei* and derivatives, including 2T1 [17], were cultured in HMI-11 (Gibco) supplemented with 10% foetal bovine serum (Sigma) in a humidified incubator at 37 °C with 5% CO_2_. Genetic manipulations were carried out by electroporation using a Nucleofector (Lonza), with cytomix for routine transfections and a Human T-cell kit (Lonza) for high efficiency library transfections [10]. Selection of recombinant bloodstream form clones was carried out by the addition of phleomycin, G418, puromycin, hygromycin and/or blasticidin at 2, 2, 2, 2.5 and 10 μg/mL, respectively; clones were subsequently maintained in 1, 1, 1, 1 and 2 μg/mL of each antibiotic, respectively. Transcription was inhibited by treating cells with 5 µg/mL of actinomycin D (Sigma). Tetracycline was used at 1 μg/mL to induce RNAi knockdown and inducible expression of Cas9. Cell density was determined using a haemocytometer.

### Plasmid construction

To generate the Neo^R^ *ZIP3* 3’-UTR construct, a PCR fragment corresponding to the full 1082 bp inter-CDS region of *ZIP3* was amplified from wild-type *T. brucei* genomic DNA using primers P9, carrying a BamHI site and P10, carrying a BglII site. The plasmid pRPa^iUTR^ [11] was digested with BamHI and BglII to remove a BSD-TK_Ald 3’-UTR fragment, and the PCR product was inserted into those sites. The recombinant plasmid was linearised with PmeI and transfected into *T. brucei*. To generate the *ZNK1* RNAi construct (pRPa^iSL^_11.9510), a 517 bp Tb927.11.9510 CDS fragment was selected using the RNAit tool [40]. This fragment was amplified by PCR from *T. brucei* genomic DNA and cloned in the tetracycline-inducible RNAi construct, pRPa^iSL^ [41]. The forward primer (P5) carried ApaI and KpnI sites, and the reverse primer (P6) carried XbaI and BamHI sites for cloning in either orientation.

The resulting pRPa^iSL^_11.9510 plasmid was verified by diagnostic restriction digestion and sequencing, then linearised with AscI prior to transfection into *T. brucei*. To generate the N-terminal 6×myc ZNK1 tagging construct, a 1134 bp fragment corresponding to the 5’ region of the Tb927.11.9510 gene was amplified by PCR from *T. brucei* genomic DNA using primers that introduced an AvrI restriction site at the 5’ end (P7) and a BamHI site at the 3’ end (P8). The PCR product was digested with AvrI and BamHI and cloned into the corresponding sites in pNAT_BSD^6xmyc^ [41]. The resulting plasmid was verified by restriction digestion and sequencing, linearised with SalI and transfected into *T. brucei*.

### RT-qPCR

Total RNA was extracted from bloodstream-form *T. brucei* cells using the RNeasy kit (Qiagen), including on-column DNase digestion, according to the manufacturer’s instructions. First-strand cDNA was synthesised from 1–2 µg of DNase-treated RNA using M-MLV™ Reverse Transcriptase (Sigma) and oligo-dT primers. Quantitative real-time PCR was performed on a QuantStudio 3 (Thermo Fisher Scientific) using a Luna^®^ Universal qPCR Master Mix (New England Biolabs) and gene-specific primers for Tb927.11.8990/*ZIP3’* (P15+P16), Tb92711.9000-9030/*ZIP3* (P17+P18), Tb927.11.9510/*ZNK1* (P19+P20), Tb927.5.4140/*ZIP4* (P21+P22), Neo^R^/*NPTII* (P23+P24), and the reference gene tubulin (P25+P26). Reactions were run in technical triplicate, and a no-RT control was included to confirm absence of genomic DNA contamination. Melting-curve analysis was used to verify amplification specificity. Relative mRNA abundance was calculated using the 2^-ΔΔCt^ method, normalising Ct values to the reference gene and expressing data relative to the control.

### Dose-response assay

*T. brucei* cells were seeded at 2 x 10^3^ cells/mL in 96-well plates containing increasing concentrations of G418 in quadruplicate. Sixty-six hours later, 20 µL of 0.125 mg/mL resazurin was added. Plates were incubated for a further 6 h prior to measuring cell viability using a plate reader with the following settings: λexc 530 nm and λem 585 nm. Data were converted to “% Survival” by subtracting experimental values from the average of HMI-11 (media)-only control, followed by calculation of proportion of *T. brucei*-only control. The half-maximal effective concentration (EC_50_) was derived from nonlinear regression analysis, using the “Sigmoidal dose-response (variable slope)” function in GraphPad Prism 8 (GraphPad Software, Inc., La Jolla, CA, USA).

### RNAi Target sequencing (RIT-Seq)

The RIT-Seq screen was carried out essentially as described [10,42], except that cells were selected in 8 μg/ml G418 for seven or ten days prior to extraction of genomic DNA. High-throughput sequencing was on a DNBseq platform (Illumina) at the Beijing Genomics Institute (BGI). The sequencing data analysis pipeline was adapted from [43]. The FASTQ files with forward and reverse paired end reads (4 technical replicates for each samples) were concatenated and aligned to the reference genome v46 of *T. brucei* clone TREU927 downloaded from TriTrypDB [9] using Bowtie2 (2.3.5) [44] with the ‘very-sensitive-local’ pre-set alignment option. The alignments were converted to BAM format, reference sorted and indexed with SAMtools (1.9) [45]. The alignments were deduplicated with the Picard tools package using the MarkDuplicates function (http://broadinstitute.github.io/picard/); to minimise the potential for overrepresentation of the shortest RIT-seq fragments. Alignments with properly paired reads were extracted with SAMtool view using the -f 2 option and parsed with a custom python script to extract the paired reads containing the index sequence (GTGAGGCCTCGCGA) in forward or reverse complement orientation. The genome coverage of the aligned reads was extracted from the bam files using deeptools (3.5) in bedGraph format [46]. The bedGraph read-mapping files were visualised with the svist4get python package [47].

### *In silico* analysis of motif-enrichment in the *ZIP3* 3’-UTR

Motif enrichment analysis was performed by Multiple Em for Motif Elicitation (MEME) [48] using sequences 1 kb downstream from the stop codons of *ZIP3* genes from multiple trypanosomatids (Supplementary Data 1); one orthologue of Tb927.11.9000 from each species (data from tritrypdb.org) [9]. MEME analyses were run to identify any number of 8 b motif repetitions in the given strand only, with otherwise default conditions in classic mode. The most enriched GUGUGUG(U/C) motif was then used to determine motif frequency across the ‘427’ transcriptome (*T. brucei* Lister 427_2018 v68, from TryTripDB) using Find Individual Motif Occurrences (FIMO) analyses [49], searching the given strand only and considering a p-value <3e^-5^ (score >14). The motifs were then clustered into blocks at least 50-bp apart. The results were visualised using the ggplot2 package (version 4.0.0; https://ggplot2.tidyverse.org) and the circlize package (version 0.4.16).

### Immunofluorescence microscopy

Immunofluorescence microscopy was conducted on fixed cells in 1% formaldehyde, dried onto slides in 1% (w/v) BSA and permeabilised with 0.5% (v/v) Triton X-100. Primary mouse α-cmyc 9B11 (Cell Signalling technology) antibody was used at 1:5000 dilution, and secondary α-mouse Alexa 488 (Invitrogen) antibody was used at 1:2000. Cells were mounted in Vectashield (Vector Laboratories) containing DAPI (4′,6-diamidino-2-phenylindole). Images were acquired using a AxioImager Z1 microscope (Carl Zeiss) and analysed with ImageJ.

### Protein blotting

Cells for western blotting analysis were lysed in 4% (w/v) SDS/50 mM Tris-HCl pH 7.4 without heating, and extracts were run in 12% SDS-PAGE gels. Western blotting was performed according to standard protocols, with primary rabbit α-NPTII (Fitzgerald Industries International) and mouse α-EF1α CBP-KK1 (Merck Millipore) used at 1:5000 dilution. Detection was performed with the BIORAD Clarity^TM^ chemiluminescence kit (BioRad) using a Chemidoc (BioRad). Images were analysed using ImageLab (Biorad).

### CRISPR-Cas9-mediated gene knockout

CRISPR–Cas9-mediated disruption of ZNK1 was performed essentially as described [50]. Briefly, we designed a 20-nt single-guide RNA (sgRNA) targeting the *ZNK1* coding sequence. Complementary oligonucleotides (P11/P12) were annealed and ligated into the pT7sgRNA vector. The resulting plasmid (pT7sgRNA_11.9510) was verified by sequencing, linearised with NotI, and transfected into 2T1^T7-Cas9^ cells that express tetracycline-inducible Cas9. Cas9 expression was induced in the resulting clones, which were then transfected with an NPTII gene cassette flanked by sequences homologous to the *ZNK1* locus, using primers P13+P14. Edited clones were recovered using G418 selection. Precise *ZNK1* knockout was confirmed by diagnostic PCR and whole genome sequencing.

### Whole genome sequencing and analysis

*T. brucei* genomic DNA was extracted using DNazol (ThermoFisher), following the manufacturer’s instructions. Genome sequencing for *ZNK1* null and control strains was performed on a DNBseq platform (BGI); 3.5 Gbp/sample, 150 bp read length. Initial quality control of fastq files was performed using FastQC (https://www.bioinformatics.babraham.ac.uk/projects/fastqc/), followed by adapter trimming and quality filtering with Fastp (0.20.0) [51]. Processed reads were aligned to the Tb927 genome assembly (TriTrypDB v68) [9] using Bowtie2 (2.3.5) [44] with “– very-sensitive-local” parameters. The resulting alignments were processed with SAMtools (1.9) [45] for sorting and indexing. Bam files were transformed to bigWig track file using bamCoverage from DeepTools (3.5) [46] with –binSize 5 and – normalizeUsing RPKM. The linear coverage visualization was performed with the pyGenomeTracks [52] Python package. Bam files from parental and derived clones were analysed using bamcompare from DeepTools (3.5) [46] with –bin 200, smooth 500, –operation log2, –normalizeUsing RPKM –extendReads, –scaleFactorsMethod None, and –outFileFormat bedgraph. The circular fold-change visualization of the bedgraph files was performed with the pyCirclize (1.6) Python package (https://github.com/moshi4/pyCirclize).

### Transcriptomics

RNA-seq analysis was conducted on *ZNK1* null and control strains. Total RNA was extracted using a Qiagen RNeasy kit in triplicate according to the manufacturer’s instructions. Sequencing was performed on a HiSeq platform (Ilumina) at the Beijing Genomics Institute. Initial quality control of fastq files was performed using FastQC, followed by adapter trimming and quality filtering with Fastp (0.20.0) [51]. Processed reads were aligned to the reference TREU927 genome or Lister 427_2018 genome (TriTrypDB v68) [9] using Bowtie2 (2.3.5) [44] with ’--very-sensitive-local’ parameters. The resulting alignments were processed with SAMtools (1.9) [45] for sorting and indexing, and PCR duplicates were marked using Picard MarkDuplicates (2.22.3) [53]. Read counts per coding sequence were quantified using featureCounts (1.6.4) [54] with parameters accounting for multi-mapping reads (-M) and overlapping features (-O) and configured to count only reads where both ends were mapped (-B) and to exclude chimeric reads that mapped to different chromosomes (-C). The datasets were then used as input for differential expression analysis using the edgeR package (3.28) [55] in R (3.6.1). We retained only genes that had a minimum count of 10 reads in at least one sample and a minimum total count of 30 reads across all samples. The cqn package (1.32) [56] was used to compute gene length and GC content bias. The resulting offsets were incorporated into the DGEList object before computing dispersions. We fitted our data to a generalised linear model (GLM) using the quasi-likelihood (QL) method via glmQLFit and performed differential expression testing using the quasi-likelihood F-test through the glmQLFTest function. The output from this statistical test was then processed using the topTags function, which extracted the complete set of test results. We configured topTags to return all tested genes (n=Inf) without applying any sorting (sort.by=“none”), ensuring that the original gene order was preserved in the output. For multiple testing correction, we employed the Benjamini-Hochberg (BH) procedure (adjust.method=“BH”) to control the false discovery rate.

### Oligonucleotides

Details of oligonucleotides used for plasmid construction, RT-qPCR and ZNK1 knockout can be found in Supplementary Data 1.

## Data availability

High-throughput sequencing data; RIT-seq, genome sequencing and RNA-seq, have been deposited in the Sequence Read Archive (SRA) with BioProject ID: PRJNA1405336 (https://www.ncbi.nlm.nih.gov/bioproject/PRJNA1405336).

## Author Contributions

The experiments were designed by T.L., A.T., A.M.T. and D.H. and carried out by T.L., A.T., M.D., and I.J.V. High-throughput sequencing data analysis was performed by M.T. *Cis*-regulatory motif searching and analysis was performed by G.B-R. The work was supervised by L.M.F., A.M.T. and D.H. The manuscript was written by T.L., A.M.T. and D.H. The manuscript was edited by all authors.

## Competing interests

The authors declare no competing interests.

Supplementary Information is available for this paper.

Correspondence and requests for materials should be addressed to

---

**Supplementary Figure 1.**
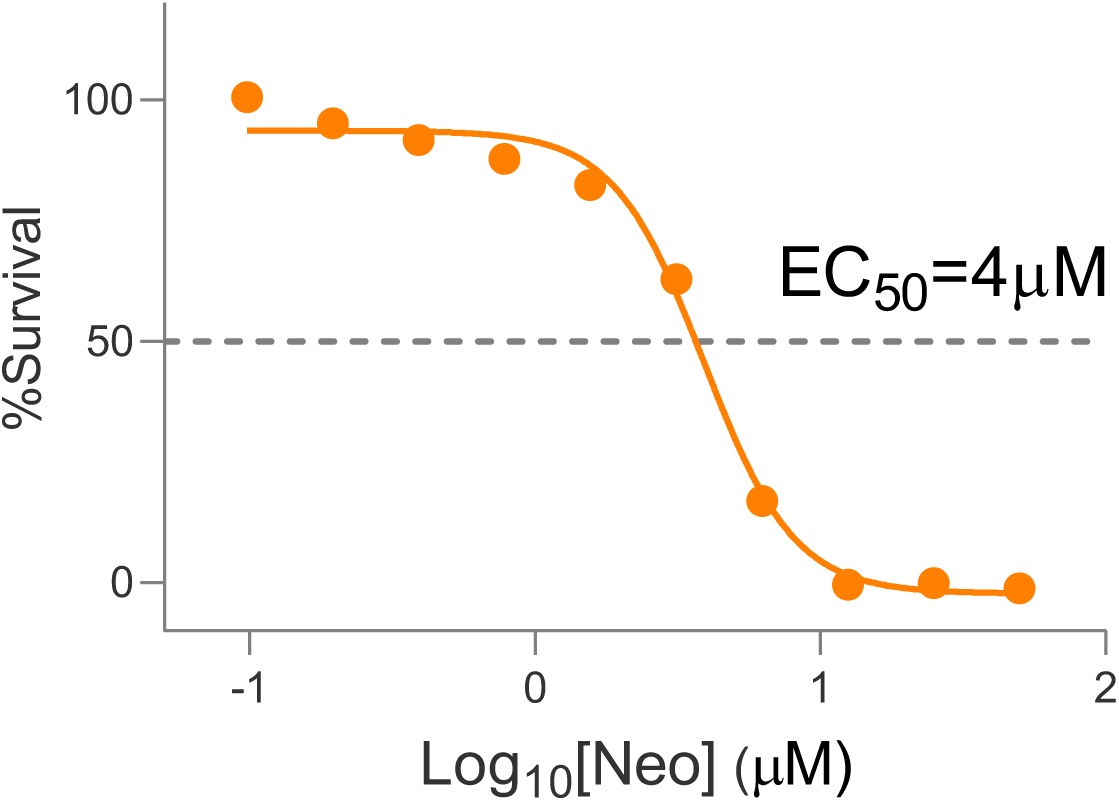
Dose-response curve for the *T. brucei* Neo^R^ *ZIP3* 3*’*-UTR reporter strain in G418. We used 2 x EC_50_ for RIT-seq library screening to select for loss of *ZIP3* 3*’*-UTR-based negative control.

**Supplementary Figure 2.**
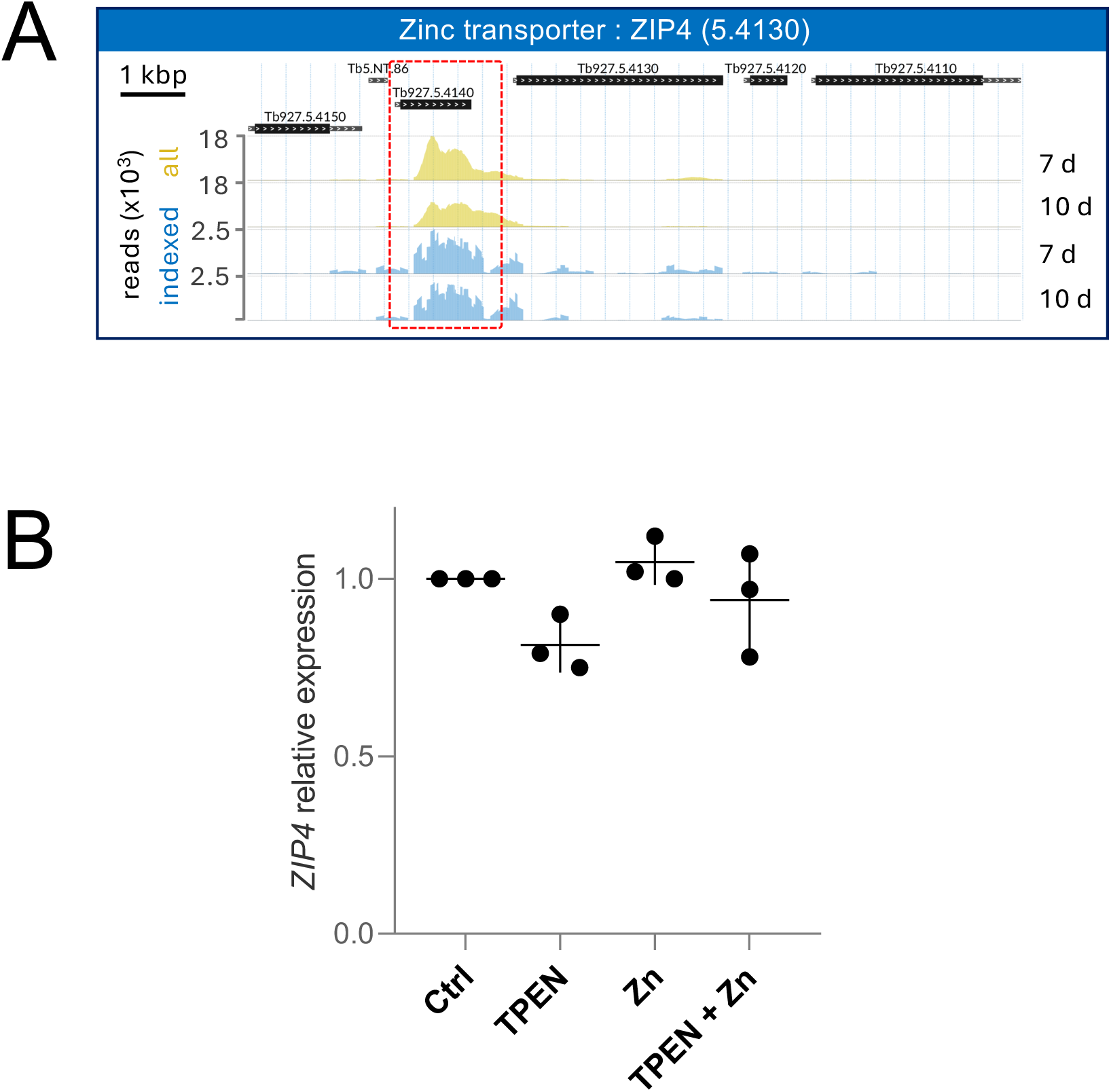
RIT-seq identifies *ZIP4* (Tb927.5.4140) as a hit. (**A**) The profile shows enrichment of RIT-seq fragments at the *ZIP4* (Tb927.5.4140) locus. Other details as in Fig. 2D. (**B**) RT-qPCR assays were used to assess the abundance of *ZIP4* transcripts in *T. brucei* grown in zinc-sufficient or zinc-limiting conditions. Other details as in Fig. 1A.

**Supplementary Figure 3.**
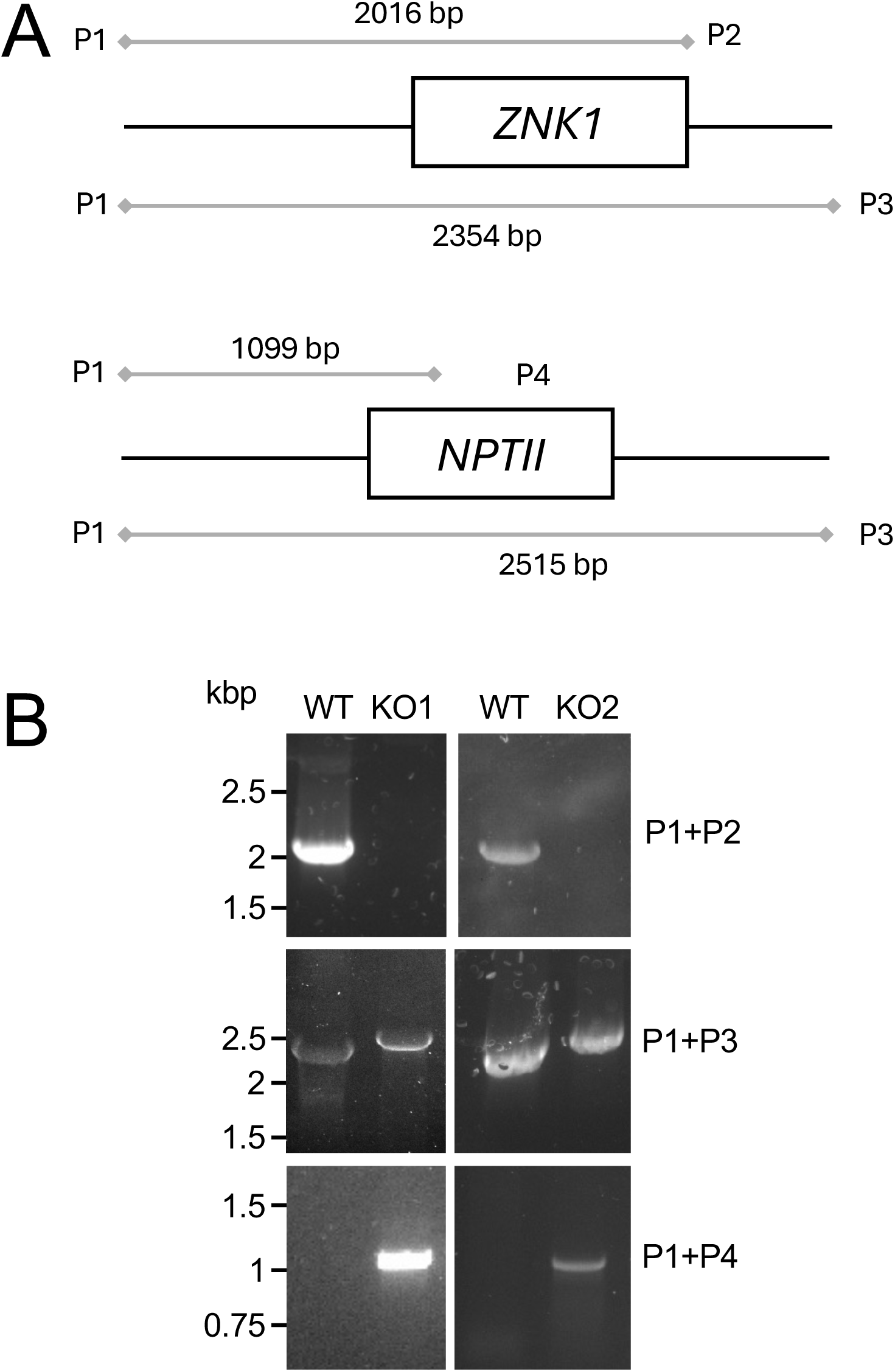
Knockout of *T. brucei ZNK1* (**A**) The schematic maps show the *ZNK1* locus before (upper panel) and after (lower panel) CRISPR-Cas9-mediated knockout. Primers used for the PCR assays are shown. (**B**) PCR assays using genomic DNA before and after *ZNK1* knockout; two independent knockout strains were analysed (KO1 and KO2).

**Supplementary Figure 4.**
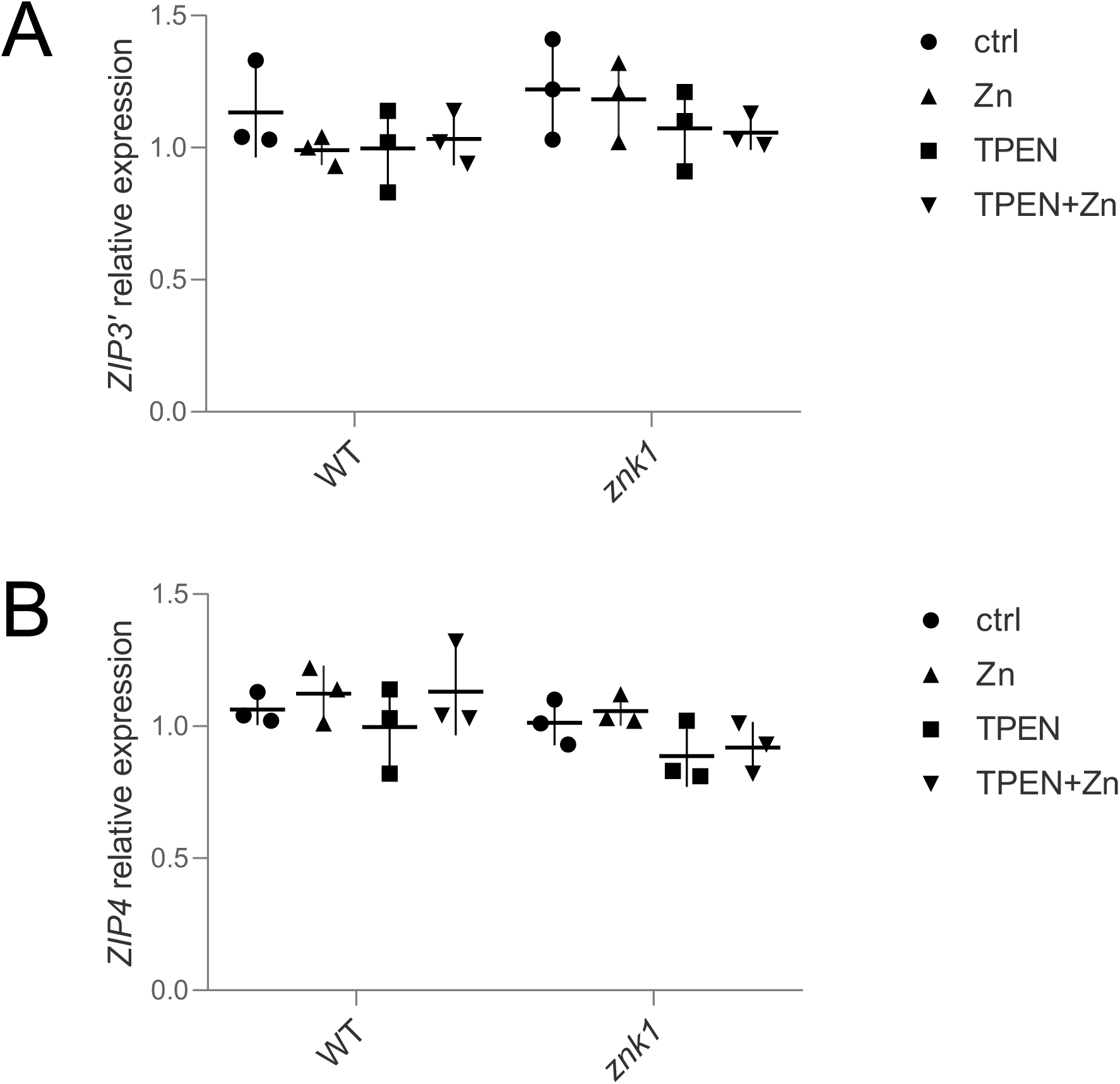
RT-qPCR assays were used to assess the abundance of *ZIP3’* (**A**) or *ZIP4* (**B**) transcripts in wild-type or *znk1*-null *T. brucei* grown in zinc-sufficient or zinc-limiting conditions. Other details as in Fig. 1A.

